# Low serine–glycine–one-carbon pathway activity defines purine salvage dependence in neuroblastoma

**DOI:** 10.64898/2026.11.18.767701

**Authors:** Jane Ding, Thi Thinh Nguyen, Mohit Bansal, Sunil Sudarshan, Yanfeng Zhang, Han-Fei Ding

**Affiliations:** Department of Pathology; O’Neal Cancer Center; Department of Urology; Department of Genetics and Precision Medicine Institute, Heersink School of Medicine, University of Alabama at Birmingham, Birmingham, AL 35294, USA

## Abstract

MYCN amplification promotes serine–glycine–one-carbon (SGOC) metabolism in neuroblastoma, creating an intrinsic difference in SGOC activity between MYCN-amplified and non-MYCN-amplified high-risk tumors. This metabolic distinction provides a model to investigate how SGOC activity shapes purine metabolism. Here, we show that non-MYCN-amplified neuroblastoma cell lines exhibit lower SGOC pathway activity and are more resistant to pharmacologic suppression of SGOC metabolism by SHMT inhibition under serine- and glycine-deprived conditions. Stable-isotope tracing revealed that non-MYCN-amplified cells exhibited lower contribution of serine-derived glycine and one-carbon units to de novo purine synthesis but elevated basal hypoxanthine and adenine salvage. These cells also showed greater preservation of adenylate pools during SHMT inhibition. Genetic depletion of APRT or HPRT1 preferentially impaired proliferation of non-MYCN-amplified cells, demonstrating increased salvage dependence. SHMT inhibition increased purine salvage, providing additional evidence that suppression of SGOC metabolism shifts purine production toward salvage. We also identify methionine salvage as an endogenous source of adenine that supports proliferation during SHMT inhibition through an MTAP–adenine–APRT axis. Together, these findings establish SGOC pathway activity as an intrinsic determinant of whether cells preferentially rely on de novo purine synthesis or purine salvage and identify salvage dependence as a potential therapeutic vulnerability of low-SGOC tumors.

## Introduction

Neuroblastoma is a biologically heterogeneous pediatric malignancy in which amplification of the MYCN oncogene defines a major subset of high-risk disease ^1–3^. MYCN drives extensive metabolic reprogramming to support the biosynthetic demands of rapid tumor growth ^4^. However, a substantial fraction of high-risk neuroblastomas lack MYCN amplification and remain clinically aggressive, indicating that alternative metabolic programs can sustain tumor growth in the absence of MYCN amplification. In previous work ^5^, we demonstrated that MYCN directly activates the serine–glycine–one-carbon (SGOC) metabolic pathway and renders MYCN-amplified neuroblastoma cells highly dependent on SGOC activity for proliferation. These findings established SGOC metabolism as an important metabolic vulnerability of MYCN-driven neuroblastoma but also raised the question of how high-risk neuroblastomas lacking MYCN amplification sustain tumor growth with lower SGOC pathway activity.

Purine nucleotides can be generated through two principal routes: de novo synthesis and purine salvage. De novo purine synthesis constructs the purine ring through a multistep pathway requiring glycine, glutamine, aspartate, and two one-carbon units supplied by 10-formyl-tetrahydrofolate, thereby creating a direct metabolic connection between SGOC activity and purine production ^6,7^. In contrast, purine salvage recycles preformed nucleobases through adenine phosphoribosyltransferase (APRT) and hypoxanthine–guanine phosphoribosyltransferase (HPRT1), providing an alternative and more energy-efficient route to nucleotide production ^8,9^. Recent isotope-tracing studies have demonstrated that salvage can contribute substantially to tumor purine pools and support tumor growth ^10,11^. However, the intrinsic metabolic determinants that govern the relative reliance of cancer cells on de novo purine synthesis versus purine salvage remain poorly understood.

Neuroblastoma provides a useful model to address this question because MYCN amplification creates intrinsic differences in SGOC pathway activity. We therefore investigated whether these differences shape purine metabolic pathway utilization. By integrating stable-isotope tracing with genetic and pharmacologic perturbation, we show that low-SGOC neuroblastoma cells exhibit elevated basal purine salvage flux and increased dependence on APRT- and HPRT1-mediated salvage, whereas high-SGOC cells rely preferentially on SGOC-supported de novo purine synthesis. We further identify methionine salvage as an endogenous source of adenine that feeds APRT-mediated purine salvage when SGOC-dependent de novo synthesis is limited. Together, these findings establish SGOC pathway activity as an intrinsic determinant of whether cells preferentially rely on de novo purine synthesis or purine salvage.

## Results

### MYCN-amplified and non-MYCN-amplified high-risk neuroblastoma tumors display distinct metabolic pathway profiles

MYCN-amplified and non-MYCN-amplified (hereafter, non-MYCN) neuroblastoma cell lines displayed comparable proliferative capacity across an expanded panel of cell lines, as assessed by doubling time (**Figure S1A**). We also observed comparable event-free survival between patients with MYCN-amplified and non-MYCN high-risk neuroblastomas in the SEQC cohort ^12,13^ (**Figure S1B**).

These findings prompted us to ask whether MYCN-amplified and non-MYCN high-risk tumors differ at the metabolic level. To gain a global view of their metabolic landscapes, we analyzed published RNA-seq datasets from two independent neuroblastoma cohorts: SEQC (total n = 498; non-MYCN high-risk n = 84; MYCN-amplified n = 92) ^12,13^ and TARGET (total n = 150; non-MYCN high-risk n = 105; MYCN-amplified n = 20) ^14^. Using single-sample GSEA (ssGSEA) ^15^, we compared metabolic pathway activity between MYCN-amplified and non-MYCN tumors. Across both datasets, 14 metabolic pathways showed consistently higher pathway-level gene expression and 15 pathways showed consistently lower pathway-level gene expression in non-MYCN tumors relative to MYCN-amplified tumors (**Figures S1C and S1D, Table S1**). Together, these analyses show that MYCN-amplified and non-MYCN high-risk neuroblastomas exhibit distinct metabolic pathway profiles.

### SGOC pathway expression and activity are lower in non-MYCN high-risk neuroblastoma

Given the central role of glycine, serine and threonine metabolism in supporting purine nucleotide synthesis through the SGOC pathway, together with the established role of MYCN in promoting SGOC metabolism ^4,5^, we focused on this pathway among those that differed between MYCN-amplified and non-MYCN tumors (**Figures S1C and S1D**). The SGOC pathway converts the glycolytic intermediate 3-phosphoglycerate into serine and glycine while generating one-carbon units that support nucleotide biosynthesis (**Figure 1A**) ^6,16,17^.

**Figure 1.**
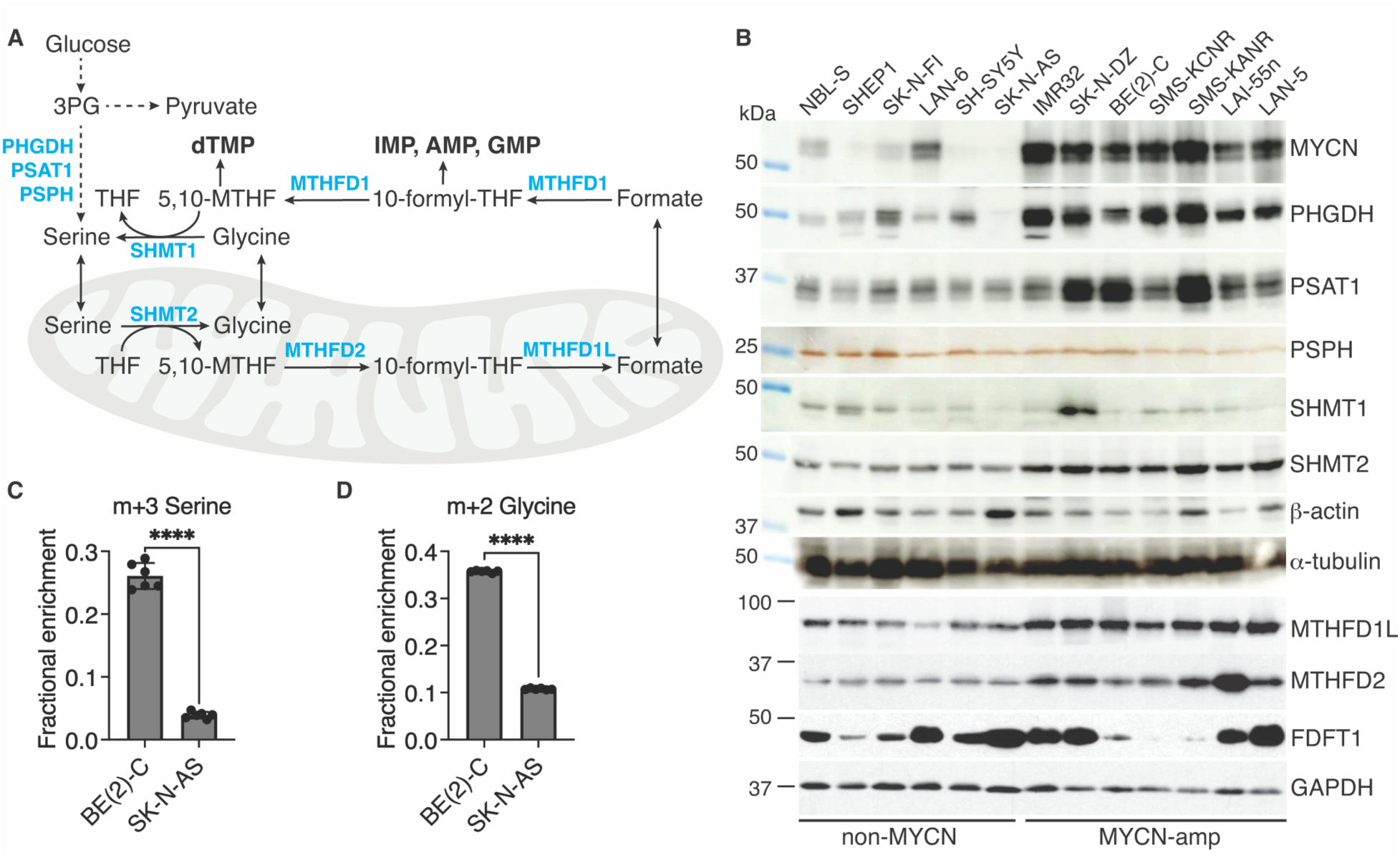
Non-MYCN neuroblastoma cells exhibit lower SGOC pathway enzyme expression and activity. (A) Schematic of the serine–glycine–one-carbon (SGOC) pathway, including conversion of the glycolytic intermediate 3-phosphoglycerate (3PG) to serine and glycine and generation of one-carbon units that support nucleotide biosynthesis. 5,10-MTHF, 5,10-methylene tetrahydrofolate; 10-formyl-THF, 10-formyl tetrahydrofolate. (B) Immunoblot analysis of SGOC pathway enzymes in MYCN-amplified and non-MYCN neuroblastoma cell lines. The cholesterol synthesis enzyme FDFT1 is shown for comparison, and β-actin, α-tubulin, and GAPDH were used as loading controls. (C-D) Fractional enrichment of serine m+3 (C) and glycine m+2 (D) following [¹³C₆]glucose tracing in MYCN-amplified BE(2)-C and non-MYCN SK-N-AS cells. Data are presented as mean ± SD (n = 6). Statistical significance was determined using an unpaired two-tailed Student’s t test. ****P < 0.0001.

Consistent with the pathway-level analysis, examination of individual SGOC gene transcripts in the SEQC cohort revealed broadly lower expression of SGOC enzymes in non-MYCN high-risk tumors compared with MYCN-amplified tumors (**Figure S2**). These included genes involved in de novo serine biosynthesis (PHGDH and PSAT1), serine–glycine interconversion (SHMT2), and downstream mitochondrial one-carbon metabolism (MTHFD1L and MTHFD2). In contrast, PSPH and SHMT1 mRNA expression did not differ significantly between MYCN-amplified and non-MYCN tumors.

To determine whether these transcriptional differences were reflected at the protein level, we examined SGOC enzyme abundance in representative MYCN-amplified and non-MYCN neuroblastoma cell lines. Consistent with the tumor transcript data, immunoblotting showed higher expression of PHGDH, PSAT1, SHMT2, MTHFD1L, and MTHFD2 in MYCN-amplified cells relative to non-MYCN cells, whereas PSPH and SHMT1 protein levels did not differ consistently between the two groups (**Figure 1B**). These results demonstrate concordant differences in SGOC enzyme expression at the transcript and protein levels according to MYCN amplification status.

To assess whether lower SGOC expression translated into diminished pathway activity, we performed stable isotope tracing using [¹³C_6_]glucose and quantified incorporation of labeled carbon into serine and glycine (**Figures 1C and 1D, Table S2**). MYCN-amplified BE(2)-C cells exhibited substantially greater fractional enrichment of serine m+3 and glycine m+2, indicating greater contribution of glucose-derived carbon to de novo serine and glycine synthesis. In contrast, non-MYCN SK-N-AS cells showed markedly lower serine m+3 and glycine m+2 enrichment, consistent with lower contribution of glucose-derived carbon to de novo serine and glycine synthesis. Collectively, these data demonstrate lower SGOC pathway expression and activity in non-MYCN neuroblastoma.

### Low-SGOC neuroblastoma cells exhibit lower serine-derived one-carbon metabolism and resistance to SHMT inhibition

Having established that non-MYCN neuroblastoma cells exhibit lower SGOC pathway expression and activity, we next asked whether this low-SGOC state is associated with altered serine-derived one-carbon metabolism. SHMT1 (cytosolic) and SHMT2 (mitochondrial) catalyze the conversion of serine to glycine while generating the one-carbon unit 5,10-methylene-tetrahydrofolate (5,10-MTHF) (**Figure 2A**). As an initial test, we examined the sensitivity of MYCN-amplified and non-MYCN neuroblastoma cell lines to the SHMT inhibitor SHIN1 ^18^ under nutrient conditions that either provide or restrict exogenous serine and glycine.

**Figure 2.**
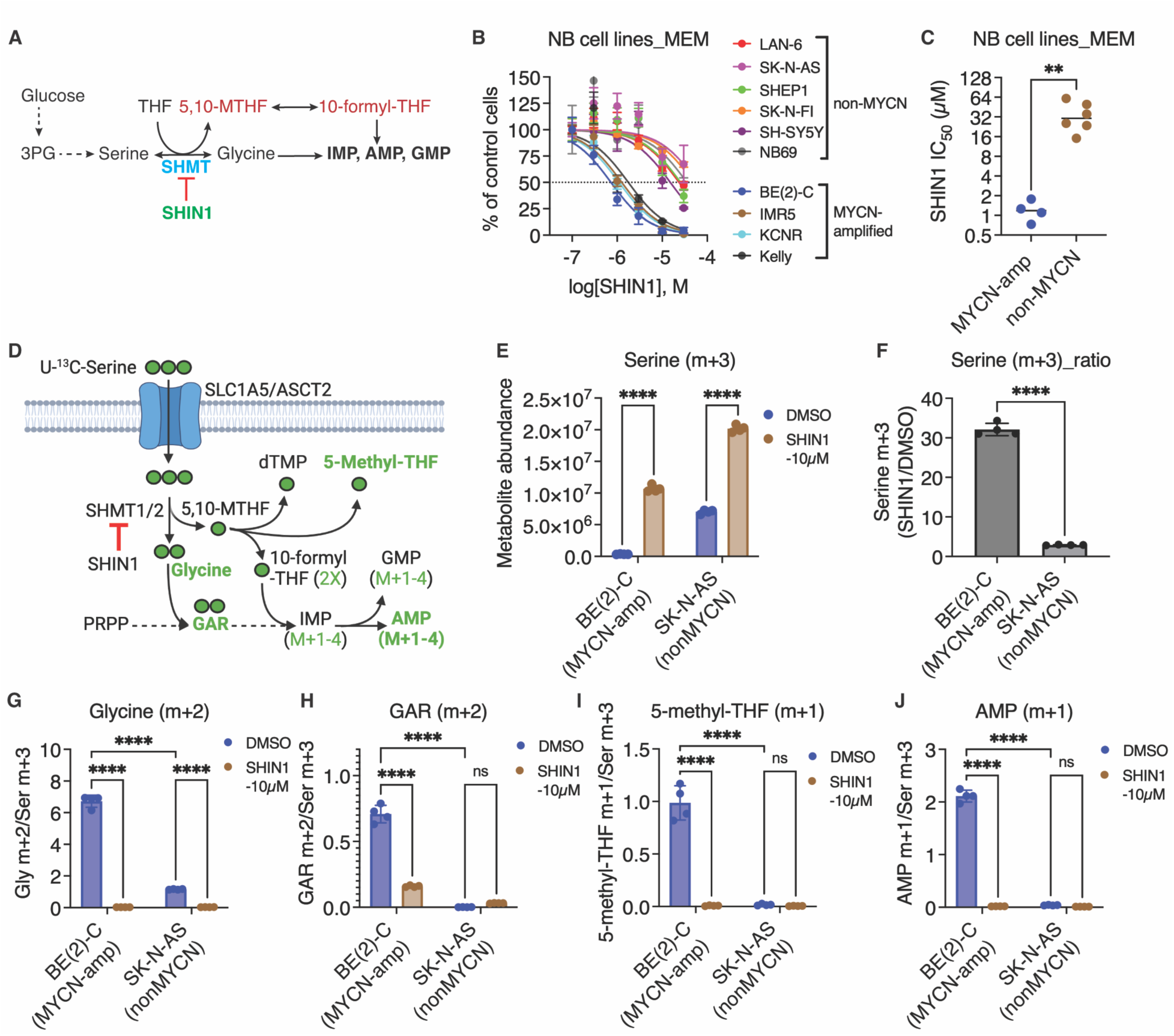
Non-MYCN neuroblastoma cells exhibit resistance to SHMT inhibition and lower serine-derived one-carbon metabolism. (A) Schematic of SHMT-mediated conversion of serine to glycine and generation of one-carbon units that support nucleotide biosynthesis. (B-C) Sensitivity of MYCN-amplified and non-MYCN neuroblastoma cell lines to the SHMT inhibitor SHIN1 in medium lacking serine and glycine, showing cell growth curves (B) and SHIN1 IC₅₀ values (C). (D) Schematic of [¹³C₃]serine tracing. (E-F) Intracellular serine m+3 abundance (E) and SHIN1-induced fold change in serine m+3 abundance (F). (G–J) Abundance of glycine m+2 (G), GAR m+2 (H), 5-methyl-THF m+1 (I), and AMP m+1 (J), normalized to intracellular serine m+3 abundance, following treatment with DMSO or SHIN1. Data are presented as mean ± SD (n = 4). Statistical significance was determined using an unpaired two-tailed Student’s t test for (C) and two-way ANOVA followed by Tukey’s multiple-comparisons test for (E–J). *P < 0.05, **P < 0.01, ***P < 0.001, ****P < 0.0001; ns, not significant.

In standard culture medium containing both serine and glycine, MYCN-amplified and non-MYCN cell lines exhibited comparable sensitivity to SHIN1, with no significant difference in IC₅₀ values between the two groups (**Figures S3A–S3B**). In contrast, when cells were cultured in medium lacking serine and glycine, which increases dependence on endogenous serine biosynthesis and SHMT-mediated generation of glycine and one-carbon units, non-MYCN neuroblastoma cell lines displayed significantly greater resistance to SHIN1 than MYCN-amplified cell lines (**Figures 2B–2C**). These findings suggest that exogenous serine and glycine can mask differences in SGOC dependence, whereas serine and glycine deprivation reveals the reduced reliance of non-MYCN cells on endogenous SGOC metabolism, as reflected by their greater resistance to SHMT inhibition.

To directly assess serine utilization and downstream one-carbon metabolism, we performed stable isotope tracing with [¹³C_3_]serine in representative MYCN-amplified BE(2)-C and non-MYCN SK-N-AS neuroblastoma cell lines in the presence or absence of SHIN1 (**Figure 2D, Table S3**). In BE(2)-C cells, intracellular m+3 serine abundance was low under control conditions but accumulated markedly (∼32.1-fold) upon SHMT inhibition (**Figures 2E–2F**). This pattern is consistent with rapid conversion of serine into downstream SHMT-dependent products under basal conditions rather than reduced serine uptake, which we previously found to be modestly higher in BE(2)-C than in SK-N-AS cells ^5^. In contrast, SK-N-AS cells showed substantially less SHMT-dependent utilization of labeled serine, as SHIN1 treatment produced only a modest (∼2.8-fold) increase in m+3 serine abundance (**Figures 2E–2F**).

Because intracellular m+3 serine abundance differed markedly between the two cell lines, downstream labeled metabolites were normalized to m+3 serine to assess the relative transfer of serine-derived carbon into downstream metabolites. By this measure, BE(2)-C cells displayed substantially greater serine-normalized labeling of downstream metabolites than SK-N-AS cells, including m+2 glycine, the purine intermediate m+2 GAR, the one-carbon metabolite m+1 5-methyl-THF, and m+1 AMP (**Figures 2G–2J**). SHIN1 markedly reduced this labeling in BE(2)-C cells. In SK-N-AS cells, serine-normalized m+2 glycine was also reduced by SHIN1, whereas GAR, 5-methyl-THF, and AMP labeling was already very low under basal conditions and showed little further reduction (**Figures 2G–2J**). These findings demonstrate lower transfer of serine-derived carbon into downstream glycine, one-carbon, and purine metabolites in non-MYCN SK-N-AS cells.

To determine whether these metabolic differences extend beyond BE(2)-C and SK-N-AS cells, we performed an independent [¹³C₃]serine tracing experiment in MYCN-amplified IMR-32 and non-MYCN SHEP1 cells (**Table S3**). SHIN1 caused substantially greater accumulation of m+3 serine in IMR-32 than in SHEP1 cells (∼11.5-fold versus ∼5.7-fold), again indicating greater basal SHMT-dependent serine utilization in MYCN-amplified cells (**Figures S3C–S3D**). Consistent with the findings in BE(2)-C and SK-N-AS cells, IMR-32 cells exhibited substantially higher serine-normalized labeling of glycine and the purine nucleotides AMP, ADP, GMP, and GTP than SHEP1 cells under basal conditions, and SHIN1 suppressed labeling of these metabolites in both cell lines (**Figures S3E–S3I**).

Together, results from two independent pairs of MYCN-amplified and non-MYCN cell lines demonstrate lower transfer of serine-derived carbon into glycine, one-carbon metabolites, and purine nucleotides in non-MYCN neuroblastoma cells, consistent with their lower dependence on SGOC metabolism.

### Low-SGOC neuroblastoma cells preserve serine, glycine, and adenylate pools upon SHMT inhibition

Having shown that low-SGOC neuroblastoma cells exhibit lower serine-derived one-carbon metabolism and resistance to SHMT inhibition, we next asked how this metabolic state affects serine, glycine, and adenylate pools during SHMT inhibition. We therefore performed targeted metabolomic profiling following SHIN1 treatment (**Table S4**). In MYCN-amplified, high-SGOC BE(2)-C cells, SHMT inhibition led to marked serine accumulation and glycine depletion (**Figures 3A–3B**), consistent with on-target blockade of SHMT-dependent serine-to-glycine conversion. Concomitantly, BE(2)-C cells exhibited pronounced depletion of the adenylate nucleotides AMP, ADP, and ATP (**Figures 3C–3E**), indicating a strong dependence on SGOC metabolism to sustain adenylate pools.

**Figure 3.**
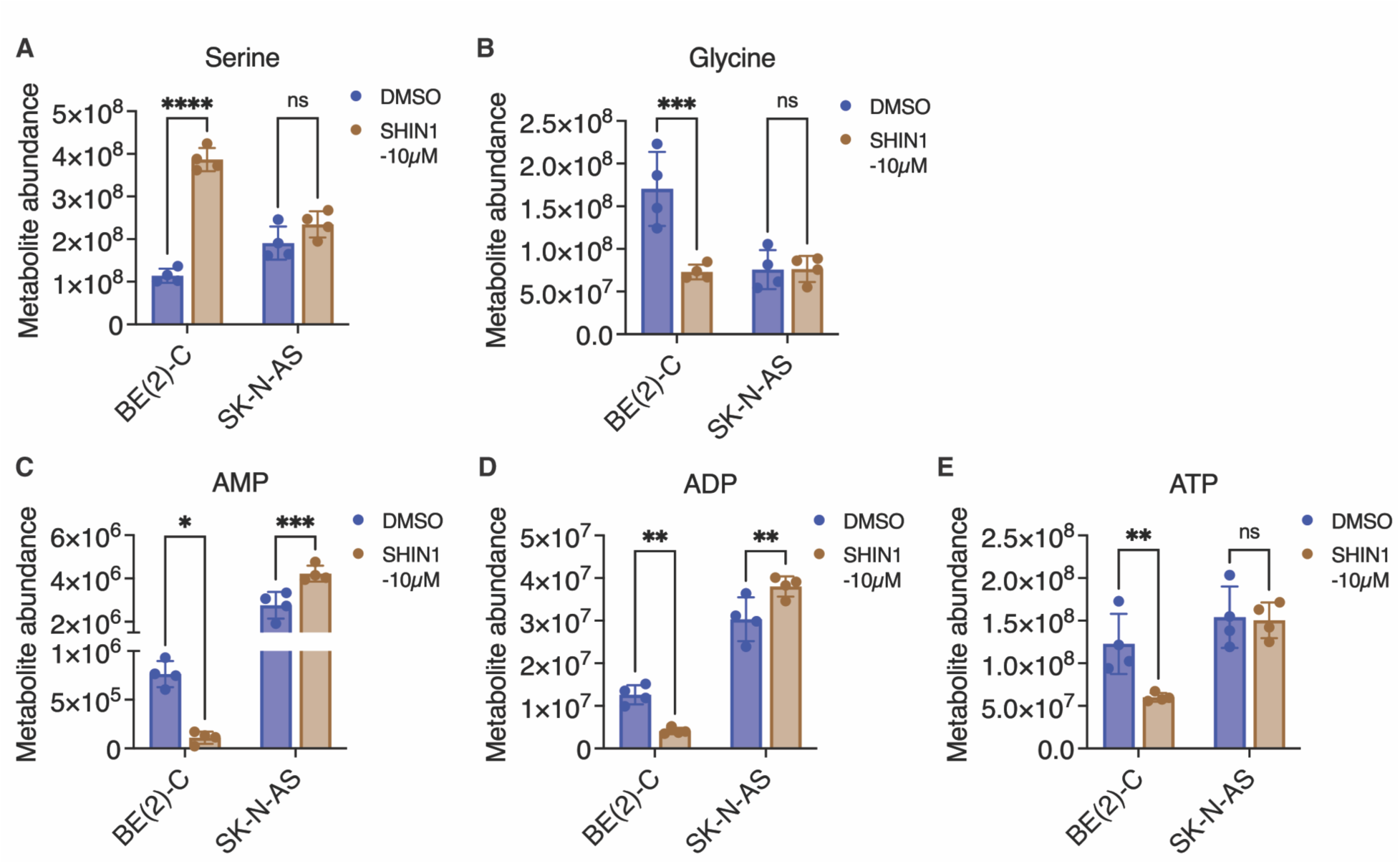
Low-SGOC neuroblastoma cells perserve serine, glycine, and adenylate pools upon SHMT inhibition Intracellular serine. (A), glycine (B), AMP (C), ADP (D), and ATP (E) abundance in MYCN-amplified BE(2)-C and non-MYCN SK-N-AS cells treated with DMSO or SHIN1. Data are presented as mean ± SD (n = 4). Statistical significance was determined by two-way ANOVA followed by Tukey’s multiple-comparisons test. *P < 0.05, **P < 0.01, ***P < 0.001, ****P < 0.0001; ns, not significant.

In contrast, SHMT inhibition had minimal effects on serine and glycine levels in non-MYCN SK-N-AS cells (**Figures 3A–3B**). Moreover, SK-N-AS cells preserved their adenylate pools, with AMP and ADP increasing modestly and ATP remaining unchanged following SHIN1 treatment (**Figures 3C–3E**). These findings indicate that non-MYCN SK-N-AS cells are substantially less dependent on SGOC metabolism to maintain adenylate pools.

Together, these results demonstrate that non-MYCN SK-N-AS cells preserve adenylate pools during SHMT inhibition, consistent with lower reliance on SGOC-dependent de novo purine synthesis.

### Low-SGOC neuroblastoma cells exhibit increased dependence on purine salvage

Having shown that low-SGOC neuroblastoma cells preserve adenylate pools during SHMT inhibition, we next asked whether purine salvage can provide an alternative route for purine production when serine-derived one-carbon metabolism is impaired. In the salvage pathway (**Figure 4A**), adenine is converted to AMP by adenine phosphoribosyltransferase (APRT), whereas hypoxanthine and guanine are converted to IMP and GMP, respectively, by hypoxanthine-guanine phosphoribosyltransferase (HPRT1). Addition of adenine or hypoxanthine markedly rescued SHIN1-induced growth inhibition in both MYCN-amplified and non-MYCN neuroblastoma cell lines (**Figures 4B–4C**), indicating that purine base salvage can support proliferation when SGOC metabolism is impaired.

**Figure 4.**
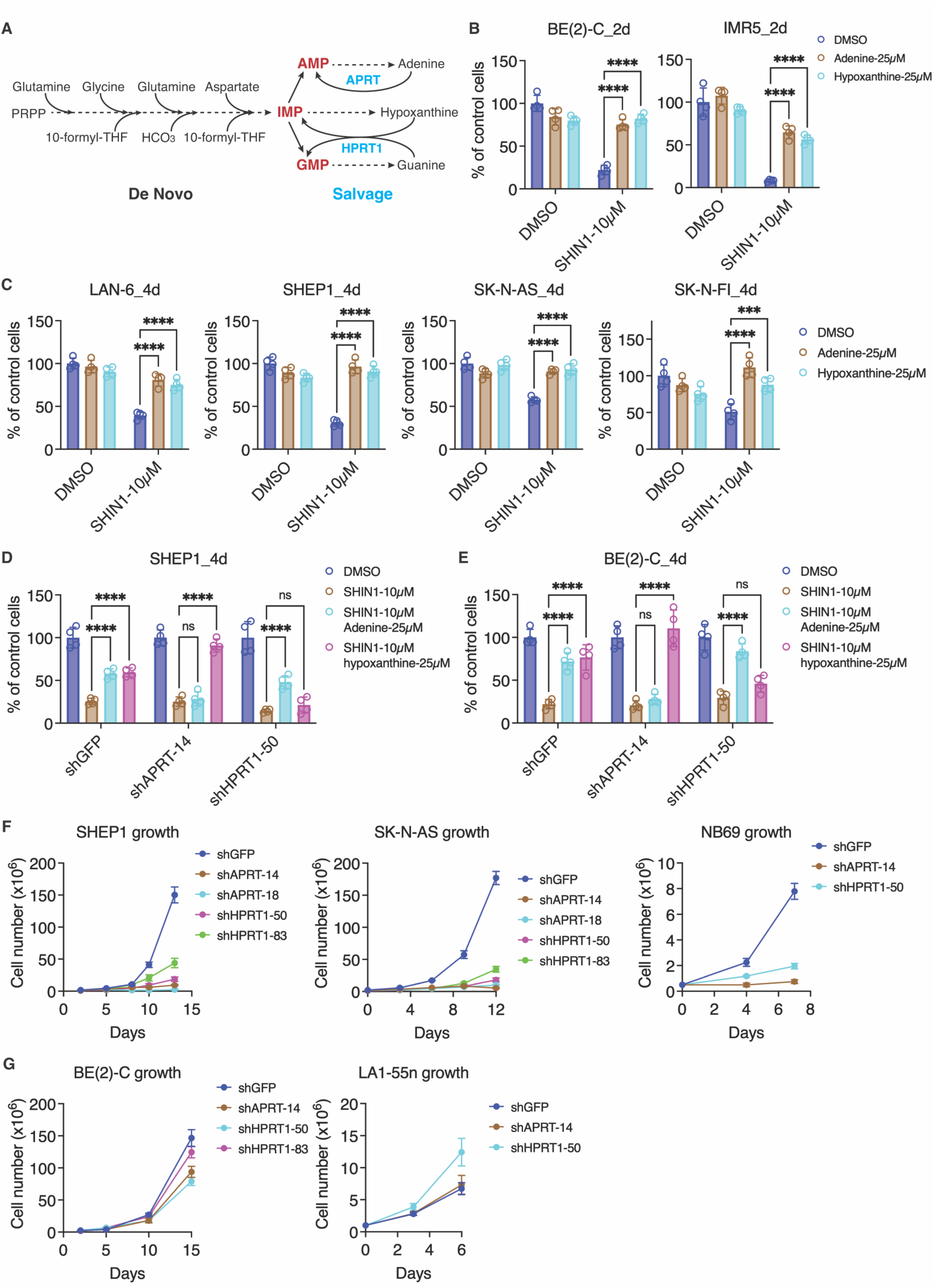
Low-SGOC neuroblastoma cells exhibit increased dependence on purine salvage. (A) Schematic of purine salvage pathways. Adenine is converted to AMP by adenine phosphoribosyltransferase (APRT), whereas hypoxanthine and guanine are converted to IMP and GMP, respectively, by hypoxanthine-guanine phosphoribosyltransferase (HPRT1). (B-C) Effect of adenine or hypoxanthine supplementation on cell viability in MYCN-amplified (B) and non-MYCN (C) neuroblastoma cell lines treated with SHIN1. (D-E) Effect of APRT or HPRT1 knockdown on adenine-or hypoxanthine-mediated rescue of SHIN1-treated cells. (F-G) Effect of APRT or HPRT1 knockdown on proliferation of non-MYCN (F) and MYCN-amplified (G) neuroblastoma cell lines. Data are presented as mean ± SD (n = 4). Statistical significance was determined by two-way ANOVA followed by Tukey’s multiple-comparisons test. ***P < 0.001, ****P < 0.0001; ns, not significant.

To determine whether this rescue requires the corresponding purine salvage enzymes, we depleted APRT or HPRT1 using independent shRNAs. Efficient knockdown of APRT and HPRT1 was confirmed at the mRNA and/or protein levels (**Figures S4A–S4C**). APRT knockdown selectively abrogated the ability of adenine to rescue SHIN1-induced growth inhibition, whereas HPRT1 knockdown specifically blocked rescue by hypoxanthine (**Figures 4D–4E**). These results demonstrate that rescue by adenine and hypoxanthine requires the corresponding APRT- and HPRT1-dependent salvage pathways.

We next asked whether MYCN-amplified and non-MYCN neuroblastoma cells differ in their dependence on purine salvage under basal conditions. Depletion of either APRT or HPRT1 markedly suppressed proliferation of non-MYCN neuroblastoma cell lines, whereas MYCN-amplified cell lines were substantially less affected (**Figures 4F–4G and S4D**). Thus, although both adenine and hypoxanthine salvage can compensate for acute SHMT inhibition, non-MYCN neuroblastoma cells are more dependent on purine salvage under basal conditions.

Together, these findings reveal increased dependence on both APRT-and HPRT1-mediated salvage in non-MYCN neuroblastoma cells and demonstrate that purine salvage can support proliferation when SGOC metabolism is impaired.

### Low-SGOC neuroblastoma cells exhibit increased purine salvage flux

Having established that low-SGOC neuroblastoma cells are more dependent on purine salvage for proliferation, we next asked whether this functional dependence is accompanied by increased salvage flux. We therefore performed stable isotope tracing with [¹⁵N₄]hypoxanthine, which is converted by HPRT1 to m+4 IMP and subsequently incorporated into adenylate and guanylate nucleotides (**Figure 5A, Table S5**).

**Figure 5.**
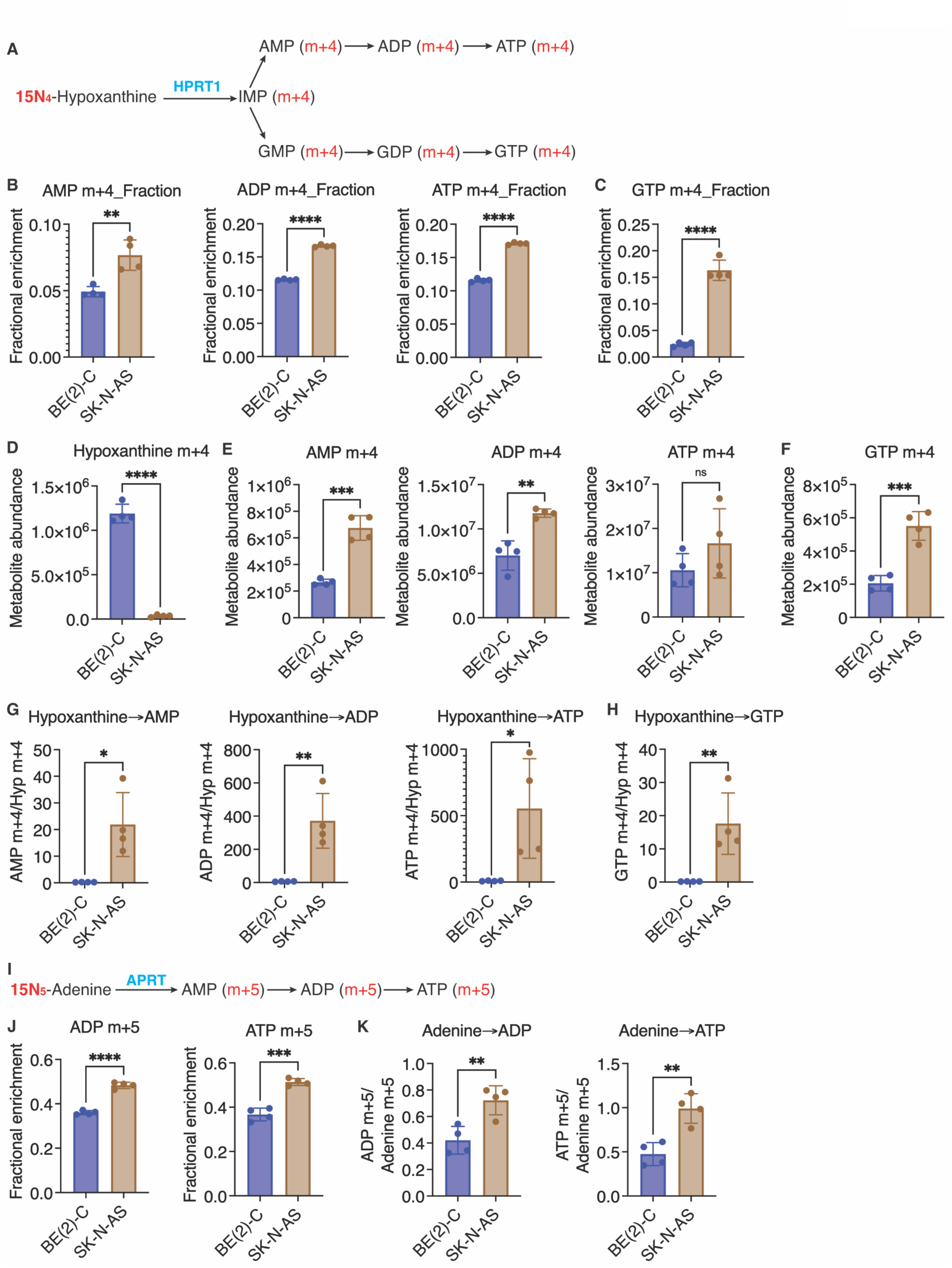
Low-SGOC neuroblastoma cells exhibit increased purine salvage flux. (A) Schematic of [¹⁵N₄]hypoxanthine salvage through HPRT1 to generate labeled IMP and downstream adenylate and guanylate nucleotides. (B-C) Fractional enrichment of m+4 AMP, ADP, and ATP (B), and GTP (C) following [¹⁵N₄]hypoxanthine tracing. (D) Intracellular abundance of m+4 hypoxanthine following tracer exposure. (E-F) Abundance of m+4 AMP, ADP, and ATP (E), and GTP (F). (G-H) Ratios of m+4 AMP, ADP, and ATP (G), and GTP (H) to intracellular m+4 hypoxanthine, shown as measures of labeled hypoxanthine utilization. (I) Schematic of [¹⁵N5]adenine salvage through APRT to generate labeled adenylate nucleotides. (J) Fractional enrichment of m+5 ADP and ATP following [¹⁵N₅]adenine tracing. (K) Ratios of m+5 ADP and ATP to intracellular m+5 adenine, shown as measures of labeled adenine utilization. Data are presented as mean ± SD (n = 4). Statistical significance was determined using unpaired two-tailed Student’s t tests. *P < 0.05, **P < 0.01, ***P < 0.001, ****P < 0.0001; ns, not significant.

We first compared hypoxanthine salvage between BE(2)-C and SK-N-AS cells under basal conditions. SK-N-AS cells exhibited significantly greater fractional enrichment of m+4 AMP, ADP, ATP, and GTP than BE(2)-C cells (**Figures 5B–5C**), indicating greater hypoxanthine salvage into purine nucleotides. Consistent with increased utilization of the labeled substrate, the intracellular m+4 hypoxanthine pool was nearly depleted in SK-N-AS cells after the tracing period but remained abundant in BE(2)-C cells (**Figure 5D**). This greater depletion of m+4 hypoxanthine was accompanied by increased accumulation of downstream labeled purine nucleotides. SK-N-AS cells contained significantly higher levels of m+4 AMP, ADP, and GTP than BE(2)-C cells, while m+4 ATP showed the same directional increase (**Figures 5E–5F**).

To further assess hypoxanthine utilization, we normalized labeled nucleotides to intracellular m+4 hypoxanthine. The ratios of m+4 AMP, ADP, ATP, and GTP to m+4 hypoxanthine were markedly higher in SK-N-AS cells than in BE(2)-C cells (**Figures 5G–5H)**, providing additional support for increased HPRT1-dependent hypoxanthine salvage in SK-N-AS cells.

To determine whether increased salvage in SK-N-AS cells extends beyond the HPRT1-dependent hypoxanthine pathway, we performed an independent tracing experiment with [¹⁵N₅]adenine, which is salvaged by APRT into adenylate nucleotides (**Figure 5I, Table S6**). Under basal conditions, SK-N-AS cells exhibited significantly greater fractional enrichment of m+5 ADP and ATP than BE(2)-C cells (**Figure 5J**). In addition, the ratios of m+5 ADP and ATP to intracellular m+5 adenine were significantly higher in SK-N-AS cells (**Figure 5K**), providing independent evidence for increased adenine salvage.

We next asked whether suppression of SGOC metabolism increases purine salvage. SHIN1 increased the fractional enrichment of m+4 ADP, ATP, and GTP in both MYCN-amplified BE(2)-C and non-MYCN SK-N-AS cells, with a similar trend observed for AMP (**Figures S5A–S5B**). indicating increased hypoxanthine salvage following SHMT inhibition. SHIN1 also increased the ratios of m+5 ADP and ATP to m+5 adenine in SK-N-AS cells, whereas no significant increase was observed in BE(2)-C cells (**Figure S5C**), suggesting that low-SGOC cells can further enhance APRT-dependent adenine salvage when SGOC metabolism is impaired.

Together, these findings demonstrate that non-MYCN SK-N-AS cells exhibit substantially higher basal purine salvage flux through both HPRT1- and APRT-dependent pathways and that SHMT inhibition can promote purine salvage. These metabolic tracing data complement the functional dependence studies and support a model in which the low-SGOC state is associated with elevated basal purine salvage flux and increased dependence on salvage pathways.

### Methionine salvage provides adenine to support purine salvage under SHMT inhibition

Having established that low-SGOC neuroblastoma cells exhibit elevated purine salvage flux and increased dependence on purine salvage for proliferation, we next sought to identify endogenous metabolic sources that could supply purine bases for salvage. [¹³C₆]glucose tracing revealed substantially greater enrichment of partially labeled methionine isotopologues in non-MYCN SK-N-AS cells than in MYCN-amplified BE(2)-C cells, particularly m+3 and m+4 (**Figure 6A, Table S2**). This labeling pattern is consistent with greater methionine salvage, in which carbons derived from the ribose moiety of methylthioadenosine (MTA) are recycled into methionine ^19–21^. Methionine salvage is coupled to polyamine biosynthesis: S-adenosylmethionine (SAM) is decarboxylated to support spermidine and spermine synthesis, generating MTA as a byproduct. MTAP cleaves MTA to generate adenine, while downstream reactions regenerate methionine and produce formate (**Figure 6B**).

**Figure 6.**
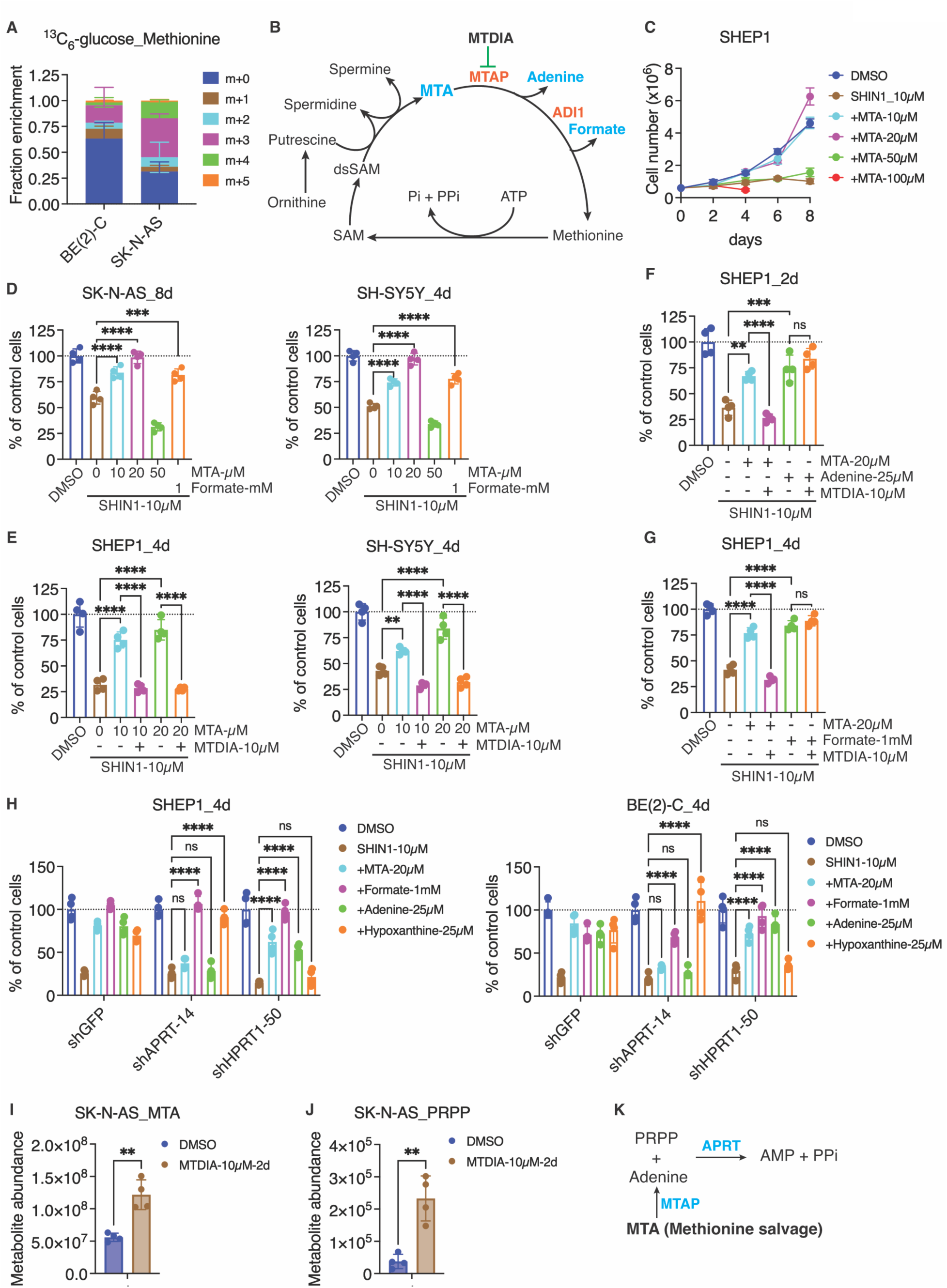
Methionine salvage supplies adenine to support purine salvage under SHMT inhibition. (A) Fractional enrichment of methionine isotopologues following [¹³C₆]glucose tracing in MYCN-amplified BE(2)-C and non-MYCN SK-N-AS cells. (B) Schematic of the methionine salvage pathway linking polyamine metabolism to generation of adenine and formate. Methylthioadenosine (MTA), produced during polyamine biosynthesis, is cleaved by MTAP to generate adenine, while downstream reactions regenerate methionine and produce formate. (C) Growth of non-MYCN SHEP1 cells treated with SHIN1 in the absence or presence of the indicated concentrations of MTA. (D) Effect of MTA or formate supplementation on growth of non-MYCN SK-N-AS and SH-SY5Y cells treated with SHIN1. (E) Effect of the MTAP inhibitor MTDIA on MTA-mediated rescue of SHIN1-treated SHEP1 and SH-SY5Y cells. (F-G) Effect of adenine (F) or formate (G) supplementation on MTDIA-mediated inhibition of MTA rescue in SHIN1-treated SHEP1 cells. (H) Effect of APRT or HPRT1 knockdown on rescue of SHIN1-treated SHEP1 and BE(2)-C cells by MTA, formate, adenine, or hypoxanthine. (I-J) Intracellular MTA (I) and PRPP (J) abundance in SK-N-AS cells following treatment with MTDIA. (K) Model illustrating the MTAP–adenine–APRT pathway through which methionine salvage supplies adenine for conversion to AMP using PRPP. Data are presented as mean ± SD (n = 6 for A; n = 4 for D–J). Statistical significance was determined by two-way ANOVA followed by Tukey’s multiple-comparisons test for (D-H) and by unpaired two-tailed Student’s t tests for (I–J). *P < 0.05, **P < 0.01, ***P < 0.001, ****P < 0.0001; ns, not significant.

To determine whether methionine salvage can support proliferation when SGOC metabolism is impaired, we examined the ability of exogenous MTA to rescue SHIN1-induced growth inhibition. MTA robustly rescued cell growth in a dose-dependent manner across multiple neuroblastoma cell lines (**Figures 6C–6D and S6A–S6B**). At higher concentrations, however, MTA reduced cell growth (**Figures 6C–6D**), likely due to feedback inhibition of polyamine synthesis ^22^. Consistent with previous reports ^18^, formate also rescued SHIN1-induced growth inhibition (**Figures 6C–6D and S6A–S6B**). These findings indicate that metabolites generated through methionine salvage can compensate for impaired SGOC metabolism.

We next determined whether MTA-mediated rescue requires MTAP, which cleaves MTA to generate adenine (**Figure 6B**). Pharmacologic inhibition of MTAP with MTDIA^23,24^ markedly impaired MTA-mediated rescue (**Figure 6E**), and MTAP knockdown similarly reduced the ability of MTA to rescue SHIN1-treated cells (**Figures S6C–S6D**). Adenine or formate bypassed the effect of MTAP inhibition (**Figures 6F–6G and S6D**), supporting an on-target effect of MTDIA.

Because MTA metabolism generates both adenine and formate (**Figure 6B**), we next asked which product is primarily responsible for MTA-mediated growth rescue. Knockdown of acireductone dioxygenase 1 (ADI1), a downstream methionine salvage enzyme involved in formate production, did not impair MTA-mediated rescue (**Figures S6D–S6G**), indicating that ADI1-dependent formate production is dispensable for this effect. In contrast, APRT knockdown largely abolished MTA-mediated rescue of SHIN1-treated cells, whereas HPRT1 knockdown did not (**Figure 6H**). These findings indicate that MTA primarily supports proliferation through MTAP-dependent adenine production followed by APRT-mediated adenine salvage.

Metabolomic analysis provided additional evidence linking methionine salvage to adenine utilization (**Table S4**). MTAP inhibition with MTDIA caused marked intracellular accumulation of MTA (**Figure 6I**), consistent with blockade of MTA metabolism. MTDIA also caused accumulation of PRPP (**Figure 6J**). Because MTAP generates adenine, which APRT converts to AMP using PRPP, the increase in PRPP is consistent with reduced PRPP utilization through the MTAP–adenine–APRT pathway. Together with the rescue and genetic studies, these findings support a model in which MTAP-dependent methionine salvage supplies adenine for APRT-mediated AMP synthesis (**Figure 6K**).

Collectively, these results identify methionine salvage as an endogenous source of adenine that supports APRT-mediated purine salvage and proliferation when SGOC metabolism is impaired.

## Discussion

Previous studies established that purine salvage can support tumor growth and becomes particularly important when de novo purine synthesis is impaired ^10,11^. However, the intrinsic metabolic factors that determine the relative reliance on de novo synthesis versus purine salvage remain poorly understood. Our study identifies SGOC pathway activity as a key determinant of this metabolic choice: high SGOC activity favors SGOC-supported de novo purine synthesis, whereas low SGOC activity shifts purine production toward greater reliance on salvage. This relationship was evident both under basal conditions and following pharmacologic suppression of SGOC metabolism by SHMT inhibition.

Several complementary observations support this model. Low-SGOC, non-MYCN neuroblastoma cells showed lower dependence on endogenous SGOC metabolism and preserved adenylate pools during SHMT inhibition, whereas high-SGOC, MYCN-amplified cells exhibited greater serine-derived contribution to glycine and purine nucleotides and were more sensitive to disruption of this pathway. Additionally, low-SGOC cells exhibited higher basal hypoxanthine and adenine salvage and greater dependence on APRT and HPRT1 for proliferation. Finally, pharmacologic suppression of SGOC metabolism by SHMT inhibition increased purine salvage. Together, these findings suggest that low SGOC activity promotes a shift from SGOC-supported de novo purine synthesis toward greater utilization of and dependence on purine salvage.

Our findings further reveal that purine salvage can be supported by an endogenous source of adenine. Metabolic tracing suggested greater methionine salvage in low-SGOC, non-MYCN SK-N-AS cells, and functional studies identified the MTAP–adenine–APRT axis as the principal route through which MTA supports proliferation of low-SGOC, non-MYCN neuroblastoma cell lines during SHMT inhibition. Because MTA is generated during polyamine synthesis, the methionine salvage pathway provides a direct metabolic connection between polyamine metabolism and purine salvage. Thus, low-SGOC cells may support their increased reliance on purine salvage not only by utilizing extracellular purine bases but also by generating adenine endogenously through methionine salvage.

These findings have therapeutic implications for high-risk neuroblastoma. MYCN-amplified neuroblastoma cells exhibit higher SGOC activity and greater dependence on SGOC-supported de novo purine synthesis, whereas non-MYCN cells exhibit lower SGOC activity, elevated purine salvage flux, and greater dependence on salvage enzymes. This metabolic divergence provides a rationale for evaluating metabolic-state-specific therapeutic strategies: targeting SGOC-dependent de novo purine synthesis may be particularly effective in MYCN-amplified, high-SGOC neuroblastomas ^5^, whereas targeting purine salvage may represent a potential vulnerability of non-MYCN, low-SGOC tumors. SGOC pathway activity may therefore provide a potential metabolic biomarker for identifying tumors preferentially dependent on de novo purine synthesis versus purine salvage.

In summary, our findings establish SGOC pathway activity as a determinant of the relative reliance on de novo purine synthesis and purine salvage. High-SGOC tumor cells preferentially rely on de novo purine synthesis, whereas low-SGOC tumor cells exhibit elevated salvage flux and increased dependence on purine salvage for proliferation. We further identify methionine salvage as an endogenous source of adenine that feeds APRT-mediated salvage, linking polyamine metabolism to purine production. Together, these findings define an intrinsic metabolic basis for purine salvage dependence and identify low-SGOC tumors as potentially vulnerable to salvage inhibition.

## Supporting information

Low SGOC_SI_Figures

Low SGOC_SI_Tables

## SUPPLEMENTAL INFORMATION

Supplemental Information includes 6 figures and 6 tables.

## ACKNOWLEDGMENTS

We thank Wilson Landon at the UAB Targeted Proteomics & Metabolomics Lab for assistance with metabolomic sample preparation and Dr. Peng Gao for assistance with stable isotope metabolomics performed at the Metabolomics Core Facility of the Robert H. Lurie Comprehensive Cancer Center at Northwestern University, supported by the NCI Cancer Center Support Grant P30 CA060553. S.S. was supported by NIH grant R01CA200653, Department of Defense grant HT94252410558, and Department of Veterans Affairs grant I01BX002930. This work was supported by NIH grants R01CA190429 and R01CA236890 and by a UAB O’Neal Invests Award to H.-F.D.

## AUTHOR CONTRIBUTIONS

J.D. and H-F.D. conceived the study and designed the experiments with contributions from T.T.N., M.B., and Y.Z. J.D., T.T.N., and H.-F.D. performed the experiments, M.B. and Y.Z. performed RNA-seq data analyses. J.D., T.T.N. and H-F.D. analyzed data with contributions from M.B. and S.S. J.D. and H-F.D. wrote the paper with contributions from Y.Z. H-F.D. supervised and provided funding for the project. All authors read the manuscript and approved its contents.

## Materials and Methods

### Cell lines and culture

Most neuroblastoma cell lines and their culture conditions have been described in detail previously^5,25,26^. IMR5, Kelly, and NB69 were gifts from J.K. Cowell (Georgia Cancer Center, Augusta University) and were cultured in RPMI 1640 (HyClone; SH30027) supplemented with 10% FBS (Atlanta Biologicals S11050). The MYCN amplification status ^27^ and neuroblastoma identity of these cell lines were routinely validated by immunoblotting and immunofluorescence for high-level nuclear expression of MYCN and the specific neuroblastoma marker PHOX2B ^28^, as appropriate. All cell lines were authenticated by short tandem repeat profiling. Cells were used within 10 passages after thawing and were free of *Mycoplasma* contamination as determined by a LookOut Mycoplasma PCR kit (Sigma-Aldrich) and staining cells with DAPI every 3 month.

Phase-contrast microscopy images of cells were captured using the EVOS M5000 Imaging System (Invitrogen). Cell proliferation was assessed using a Trypan blue assay.

### Analysis of neuroblastoma patient datasets and metabolic landscapes

Gene-expression data from two independent neuroblastoma patient cohorts, SEQC (total n = 498, GSE49711) ^12,13^ and TARGET (total n = 150) ^14^, were analyzed. For comparisons within high-risk disease, the SEQC cohort included 84 non-MYCN high-risk and 92 MYCN-amplified tumors, and the TARGET cohort included 105 non-MYCN high-risk and 20 MYCN-amplified tumors. Tumors were classified according to MYCN amplification and risk status using the accompanying clinical annotations. We retrieved 362 human pathways from the KEGG database (https://www.genome.jp/kegg/) and retained 53 metabolic pathways containing at least five annotated genes for downstream analysis. Using log2-transformed counts per million (CPM) expression values, we calculated single-sample GSEA (ssGSEA) ^15^ scores for curated metabolic pathways in individual tumors. Differences in ssGSEA scores between MYCN-amplified and non-MYCN high-risk tumors were evaluated using Wilcoxon rank-sum tests.

Pathways with a false discovery rate (FDR) < 0.05 were considered significantly different between groups, and those showing concordant differences across both cohorts were identified. All statistical analyses were conducted in R (v4.4.0) (https://www.r-project.org/).

SGOC pathway gene expression and event-free survival data for the SEQC cohort were obtained from the R2 Genomics Analysis and Visualization Platform (https://hgserver1.amc.nl/cgi-bin/r2/main.cgi). Statistical analyses and figure generation were performed using GraphPad Prism for Mac, version 11.0.2.

### Inhibitor and rescue assays

MTA (Cayman, 15593), MTDIA (Cayman, 29711), and SHIN1 (MCE, HY-112066) were dissolved in dimethyl sulfoxide (DMSO), aliquoted, and stored at −80°C until use. For SHMT inhibition assays in standard growth medium, neuroblastoma cell lines were cultured in DMEM (HyClone, SH30022) or RPMI 1640 (HyClone, SH30027), as appropriate for each cell line, supplemented with 10% FBS and treated with various concentrations of SHIN1 for 4 days. For assays in serine- and glycine-deficient medium, cells were cultured in MEM (Gibco, 11095-080) supplemented with 10% dialyzed FBS (Atlanta Biologicals, S11050), 25 mM D-(+)-glucose (Sigma-Aldrich, G8769), 1× MEM vitamins (Gibco, 11120-052), 1× sodium pyruvate (Gibco, 11360-070), 1x Glutamax (Gibco, 35050), and the non-essential amino acids alanine, asparagine, aspartic acid, glutamic acid, and proline at concentrations equivalent to those in DMEM, and treated with various concentrations of SHIN1 for 2 days.

For SHIN1 rescue assays, cells were cultured in their respective standard growth medium and treated with SHIN1 (10 µM) in the absence or presence of adenine (25 µM; Sigma-Aldrich, A2786), hypoxanthine (25 µM; Sigma-Aldrich, H9636), formate (1 mM; Sigma-Aldrich, 71539), or MTA (10–100 µM), with or without MTDIA (10 µM), as indicated.

For all inhibitor and rescue assays, cell viability was quantified by trypan blue exclusion. IC₅₀ values were calculated from fitted dose-response curves where applicable.

### Metabolomics and stable-isotope tracing

#### [^13^C_6_]glucose

[^13^C_6_]glucose tracing, metabolite extraction, instrumentation, and data processing were performed as described previously ^5^ by the West Coast Metabolomics Center, University of California, Davis. Briefly, MYCN-amplified BE(2)-C and non-MYCN SK-N-AS cells were cultured in glucose-free DMEM (Gibco, 11966025) supplemented with 10% dialyzed FBS (Hyclone, SH30079) and 20 mM [^13^C_6_]glucose (Cambridge Isotope Laboratories, CLM-1396) for 24 h. Cells were collected by scraping and centrifugation, and cell pellets were washed once with ice-cold PBS, snap-frozen in liquid nitrogen, and stored at -80°C until analysis. Six biological replicates (∼5 x 10^6^ cells per sample) were analyzed for each cell line.

#### [^13^C_3_]serine

We analyzed the contribution of serine to the production of glycine, one-carbon units, and purine nucleotides in BE(2)-C and SK-N-AS cells, with an independent validation experiment in IMR-32 and SHEP1 cells. Cells were cultured in serine-free MEM, as described above, supplemented with 280 µM [^13^C_3_]serine (Cambridge Isotope Laboratories, CLM-1574-11-0.1) and treated with DMSO or 10 μM SHIN1 for 24 h before collection for metabolite extraction.

#### [^15^N_4_]hypoxanthine and [^15^N_5_]adenine

Purine salvage was analyzed in BE(2)-C and SK-N-AS cells cultured in RPMI 1640 supplemented with 10% dialyzed FBS and pretreated with DMSO or 10 µM SHIN1 for 24h. The medium was then replaced with fresh medium containing 10 µM [^15^N_4_]hypoxanthine (Cambridge Isotope Laboratories, NLM-8500-0) or [^15^N_5_]adenine (Cayman, 41181), and cells were cultured with DMSO or 10 µM SHIN1 for 6 h before collection for metabolite extraction.

#### Steady-state metabolite profiling

BE(2)-C and SK-N-AS cells were cultured in DMEM supplemented with 10% dialyzed FBS and treated with DMSO or 10 μM SHIN1 for 24 h before collection for metabolite extraction.

#### Metabolite extraction and LC-MS analysis

For [^13^C_3_]serine, [^15^N_4_]hypoxanthine, and [^15^N_5_]adenine tracing and steady-state metabolite profiling, four biological replicates (∼5 × 10⁶ cells per sample) were analyzed for each cell line. For these experiments, culture medium was rapidly removed and cells were washed with ice-cold PBS. Intracellular metabolites were extracted with 80% methanol at −80°C, dried in a SpeedVac, and stored at −80°C until analysis. LC-MS analyses were conducted at the Metabolomics Core Facility of the Robert H. Lurie Comprehensive Cancer Center at Northwestern University. Dried metabolite extracts were reconstituted in 60% acetonitrile, vortexed, centrifuged, and the supernatants were analyzed by HPLC-MS/MS. Data were acquired and analyzed using Thermo Fisher Xcalibur 4.1 and TraceFinder 4.1, as described previously ^29^. Detected isotopologue abundances were normalized to total protein content.

### RNA interference and lentiviral transduction

Lentiviral pLKO.1 shRNA constructs were obtained from Sigma-Aldrich, including shADI1-34 (TRCN0000296534), shADI1-77 (TRCN0000052277), shAPRT-14 (TRCN0000035014), shAPRT-18 (TRCN0000035018), shHPRT1-50 (TRCN0000035050), shHPRT1-83 (TRCN0000299783), shMTAP-54 (TRCN0000256254), and shMTAP-55 (TRCN0000256255). Lentiviruses were generated in 293T Lenti-X cells (TaKaRa 632180) using the packaging plasmids pLP1 (Thermo Fisher, K497500; RRID: Addgene_22614), pLP2 (Thermo Fisher, K497500; RRID:Addgene_209989), and pLP/VSVG (Thermo Fisher, K497500, RRID:Addgene_6099). Lentiviral transductions were performed following standard protocols. Knockdown efficiency was assessed by qRT-PCR and immunoblotting.

### Quantitative real-time PCR (qRT-PCR)

Total RNA was isolated using TRIzol reagent (Thermo Fisher, 15596026). cDNA was synthesized using the iScript Advanced cDNA Synthesis Kit for RT-qPCR (Bio-Rad, 172-5038), and qRT-PCR was performed using 2X SYBR Green qPCR Master Mix (Bimake, B21203) on an iQ5 real-time PCR system (Bio-Rad, RRID:SCR_019721). The following gene-specific primers were used: APRT, forward 5’-GGCCGCATCGACTACATCG-3’ and reverse 5’-CTCAGCCTTCCCGTACTCC-3’; HPRT1, forward 5’-CCTGGCGTCGTGATTAGTGAT-3’ and reverse 5’-AGACGTTCAGTCCTGTCCATAA-3’; and B2M, forward 5’-TGCTGTCTCCATGTTTGATGTATCT-3’ and reverse 5’-TCTCTGCTCCCCACCTCTAAGT-3’. APRT and HPRT1 mRNA levels were normalized to B2M mRNA.

### Immunoblotting

Cells were lysed in SDS sample buffer and protein concentration was determined using the Bio-Rad Protein Assay Kit II (Bio-Rad, 5000002). Proteins (30-50 µg) were separated by SDS-PAGE, transferred to nitrocellulose membranes, and probed with the following primary antibodies: rabbit anti-ADI1 (1:1000, Proteintech 15133-1-AP, RRID:AB_2305014), rabbit anti-FDFT1 (1:1000, Proteintech 13128-1-AP, RRID:AB_2294094), rabbit anti-GAPDH (FL-335, 1:1000, Santa Cruz sc-25778, RRID:AB_10167668), mouse anti-HPRT (1:400, Santa Cruz sc-376938, RRID:AB_2938532), mouse anti-MTAP (1:2000, Proteintech 66706-1-Ig, RRID:AB_2882058), rabbit anti-MTHFD1L (1:1500, Proteintech 16113-1-AP, RRID:AB_2250974), rabbit anti-MTHFD2 (1:1500, Proteintech 12270-1-AP, RID:AB_2147525), mouse anti-MYCN (1:400, B8.4.B, Santa Cruz sc-53993, RRID:AB_831602), rabbit anti-PHGDH (1:300, Sigma-Aldrich HPA021241, RRID:AB_1855299), rabbit anti-PSAT1 (1:3000, Novus 21020002, RRID:AB_2172599), rabbit anti-PSPH (1:1000, E1D2X, Cell Signaling 58258), rabbit anti-SHMT1 (1:1000, D3B3J, RRID:AB_2799957), rabbit anti-SHMT2 (1:1000, Sigma HPA020549, RRID:AB_1856834), Cell Signaling 80715), rabbit anti-β-actin (1:2000, Thermo Fisher Scientific MA5-15739, RRID:AB_10979409), and mouse anti-α-tubulin (B-5-1-2, 1:5000, Sigma-Aldrich T5168, RRID:AB_477579). HRP-conjugated secondary antibodies included goat anti-mouse IgG (Jackson ImmunoResearch 115-035-146, RRID:AB_2307392) and goat anti-rabbit IgG (Jackson ImmunoResearch 111-035-046, RRID:AB_2337939). Signals were detected using Clarity Western ECL (Bio-Rad 1705061) and imaged with the Amersham ImageQuant 800 (Cytiva) or quantified using ImageJ2 (v2.16.0/1.54p, RRID:SCR_002285).

### Statistics

Quantitative data are presented as mean ± SD and were analyzed for statistical significance by unpaired, two-tailed Student’s *t*-test or two-way ANOVA followed by Tukey’s multiple-comparisons test. Unless otherwise stated, all statistical analyses were conducted using GraphPad Prism 11.0.2 for Mac (GraphPad Software, RRID:SCR_002798). *P < 0.05, **P < 0.01, ***P < 0.001, and ****P < 0.0001; ns, not significant.

