## Supplementary material for "Low serine–glycine–one-carbon pathway activity defines purine salvage dependence in neuroblastoma": Low SGOC_SI_Figures

### Inventory of Supplementary Information

#### Supplementary Figures S1-S6

Figure S1. Neuroblastoma tumor metabolic landscapes, related to Figure 1
Figure S2. SGOC gene expression in SEQC tumors, related to Figure 1
Figure S3. SHIN1 sensitivity and [$^{13}C_3$]serine tracing, related to Figure 2
Figure S4. APRT and HPRT1 knockdown validation, related to Figure 4
Figure S5. [$^{15}N_4$]hypoxanthine and [$^{15}N_5$]adenine tracing, related to Figure 5
Figure S6. Methionine salvage and MTA rescue, related to Figure 6

#### Supplementary Table S1-S6

Table S1. Metabolic landscape source data, related to Figure S1
Table S2. [$^{13}C_6$]glucose tracing data, related to Figures 1 and 6
Table S3. [$^{13}C_3$]serine tracing data, related to Figures 2 and S3
Table S4. Metabolite profiling data, related to Figures 3 and 6
Table S5. [$^{15}N_4$]hypoxanthine tracing data, related to Figures 5 and S5
Table S6. [$^{15}N_5$]adenine tracing data, related to Figures 5 and S5

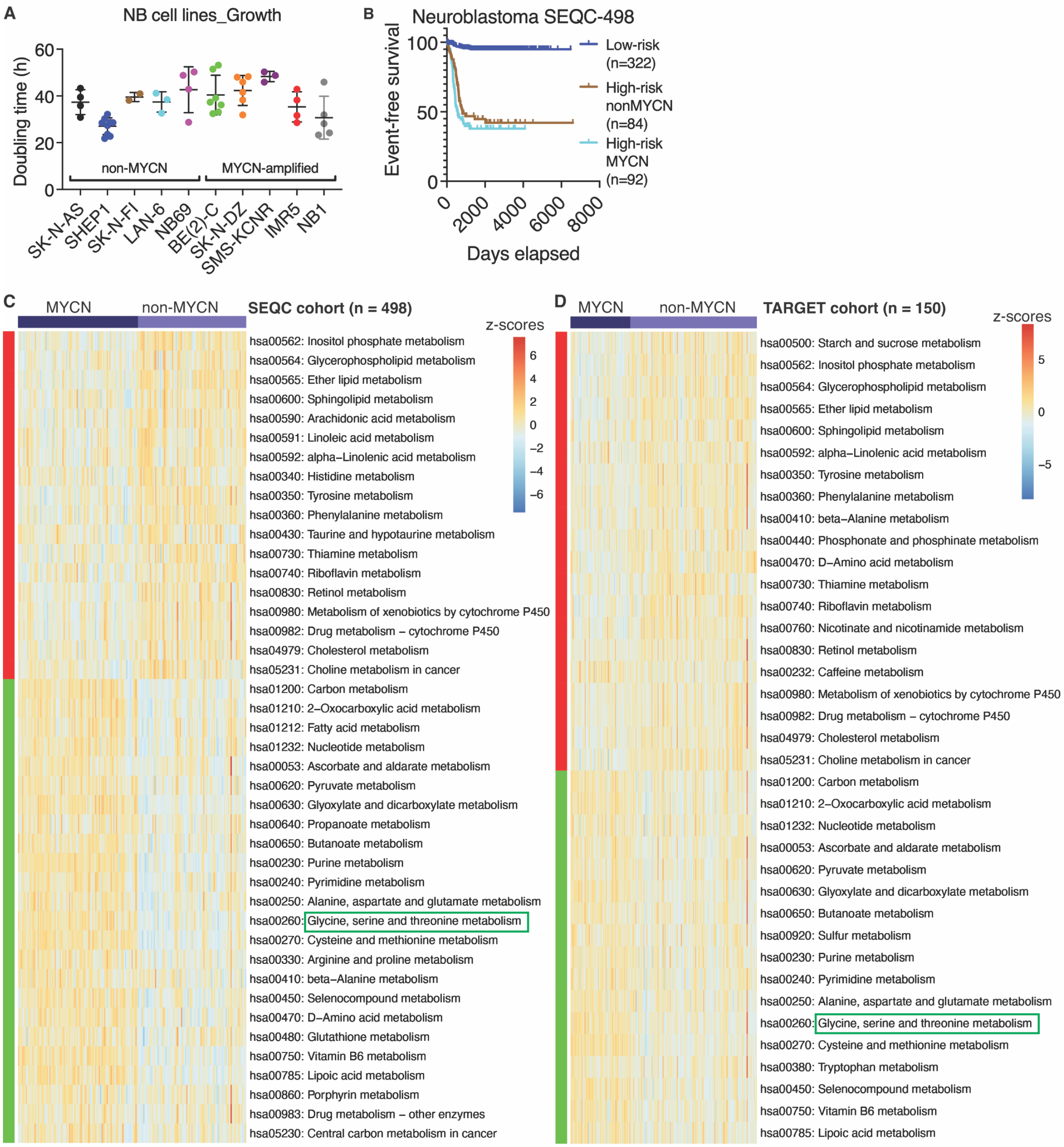


**Figure S1. MYCN-amplified and non-MYCN high-risk neuroblastomas display distinct metabolic landscapes**

(A) Cell proliferation rates of the indicated neuroblastoma cell lines as determined by cell population doubling time (DT) calculated using the formula DT = T x ln(2)/ln(Xe/Xb), where T is the incubation time in hours, Xb is the cell number at the beginning the incubation time, and Xe is the cell number at the end the incubation time.

(B) Event-free survival curves for the SEQC cohort of neuroblastoma patients.

(C–D) Heatmaps of metabolic pathways, with KEGG pathway IDs indicated, showing significant differences in metabolic pathway activity between non-MYCN and MYCN-amplified high-risk neuroblastoma tumors in the SEQC (C) and TARGET (D) cohorts (FDR < 0.05). Metabolic pathway activity was quantified using ssGSEA scores derived from mRNA expression of enzyme-encoding genes within each pathway.

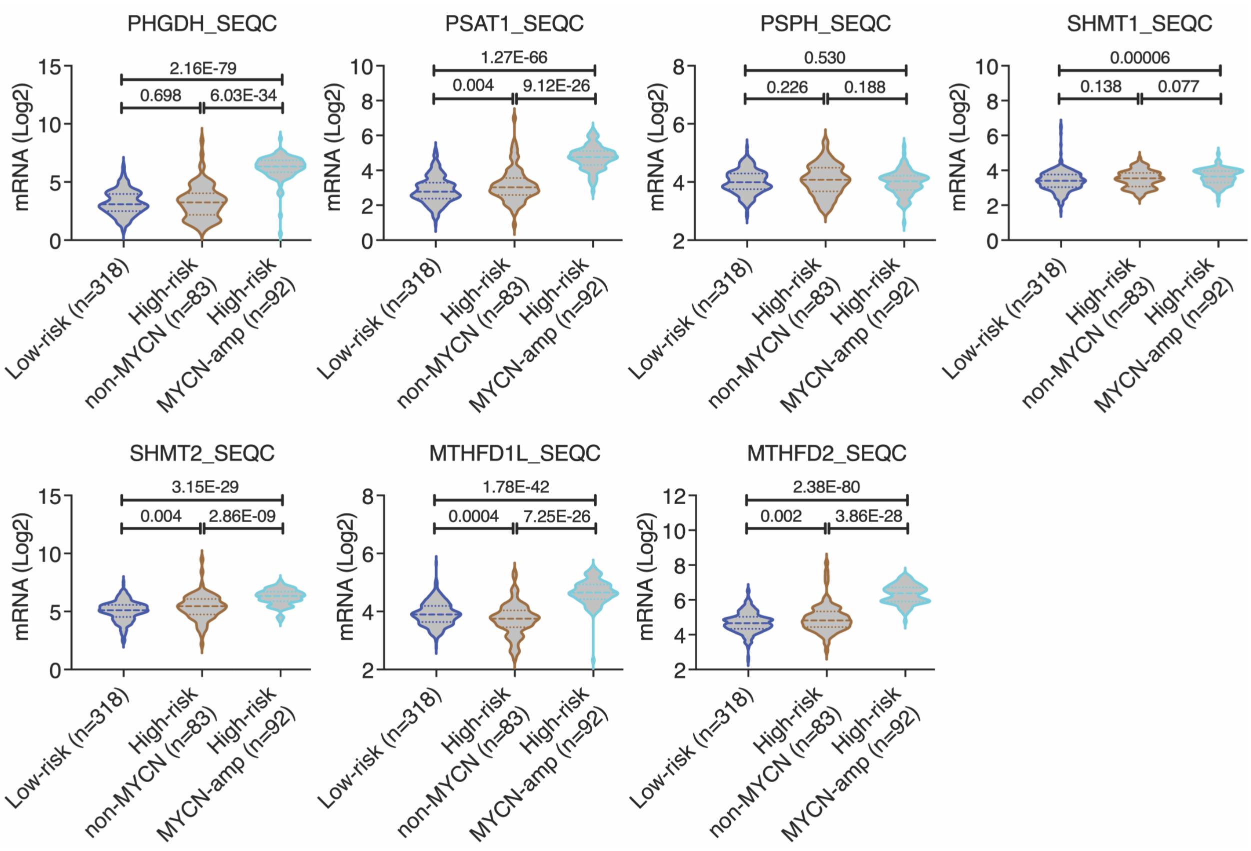


**Figure S2. SGOC pathway gene expression is lower in non-MYCN than in MYCN-amplified neuroblastomas**

Violin plots showing expression levels of individual SGOC pathway genes according to risk group and MYCN amplification status in the SEQC cohort. Data were analyzed using two-tailed Student’s t tests; P values are indicated.

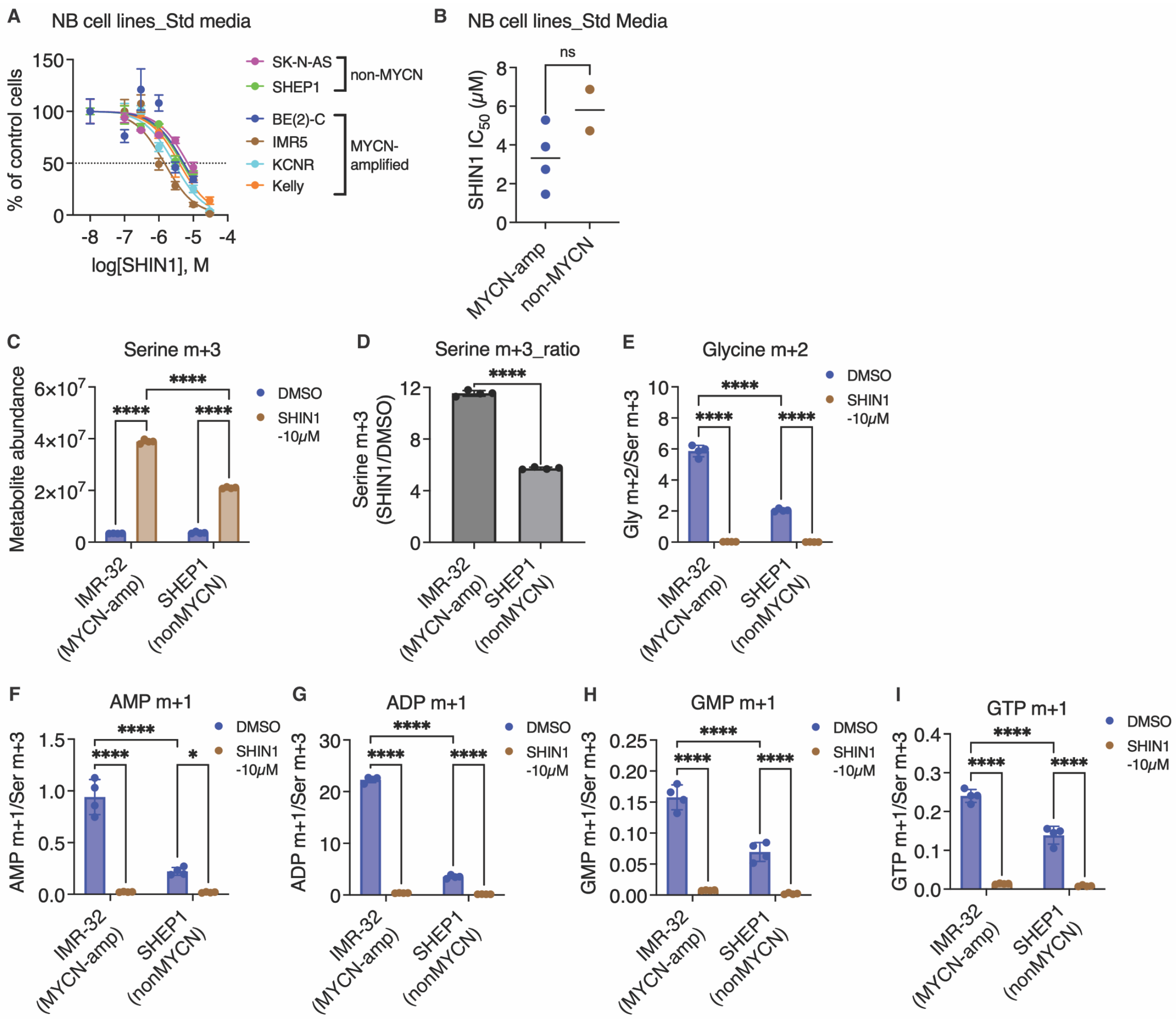


**Figure S3. SHIN1 sensitivity and [$^{13}C_3$]serine tracing in neuroblastoma cell lines**

(A) Dose-response curves for MYCN-amplified and non-MYCN neuroblastoma cell lines treated with the indicated concentrations of SHIN1 for 4 days in standard culture medium containing serine and glycine. Cell viability is shown relative to DMSO-treated controls.

(B) SHIN1 $IC_{50}$ values for MYCN-amplified and non-MYCN neuroblastoma cell lines shown in (A).

(C–I) [$^{13}C_3$]serine tracing in MYCN-amplified IMR-32 and non-MYCN SHEP1 cells, showing intracellular serine m+3 abundance (C), fold change in serine m+3 abundance following SHIN1 treatment relative to DMSO (D), and abundance of glycine m+2 (E), AMP m+1 (F), ADP m+1 (G), GMP m+1 (H), and GTP m+1 (I), normalized to intracellular serine m+3 abundance.

Data are presented as mean ± SD. Statistical significance was determined by unpaired two-tailed Student's t test for (B) and two-way ANOVA with multiple-comparisons testing for (C–I). *$P < 0.05$, ****$P < 0.0001$; ns, not significant.

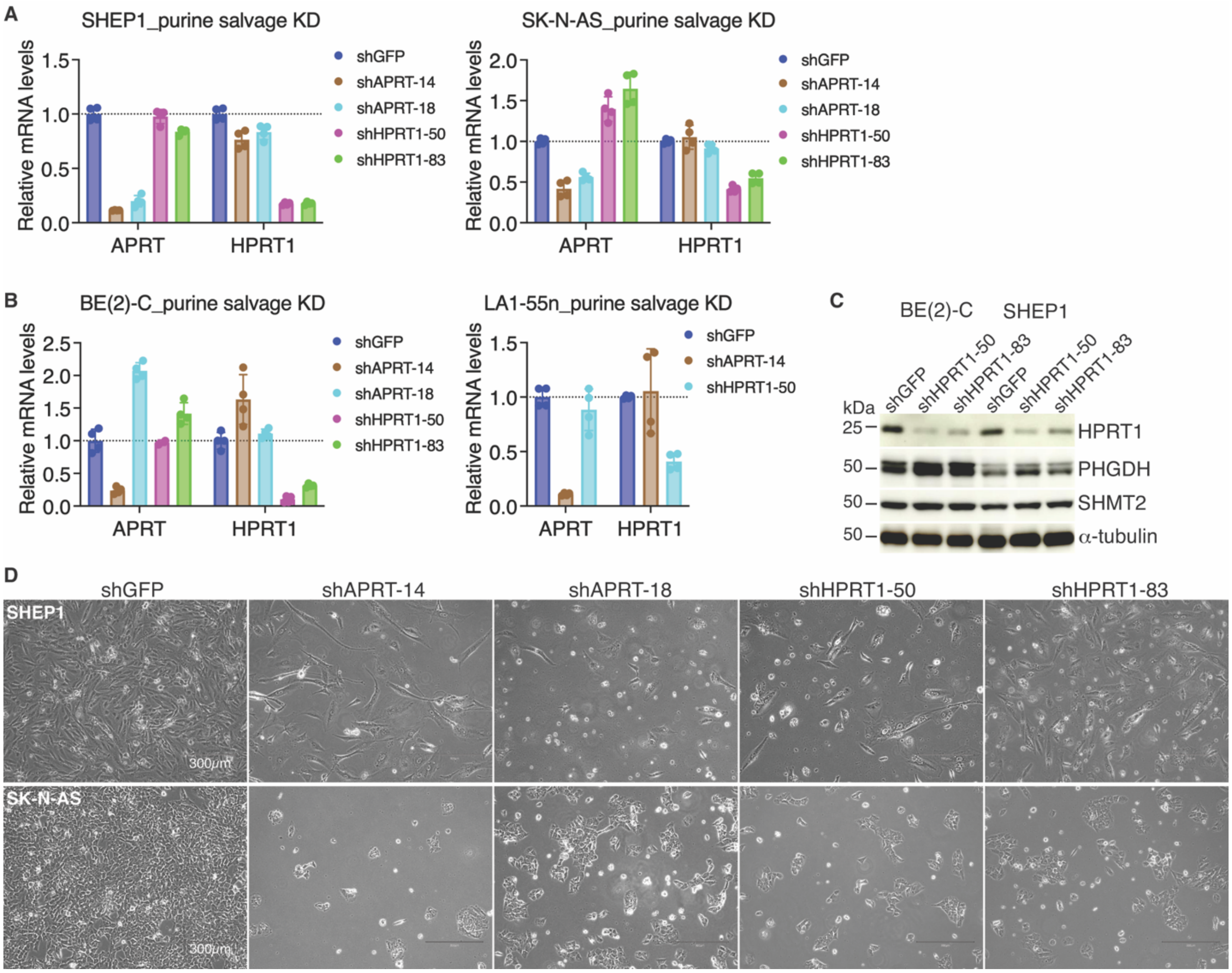


**Figure S4. Knockdown of APRT or HPRT1 inhibits growth of non-MYCN neuroblastoma cells**

(A-B) qRT-PCR analysis of APRT and HPRT1 mRNA levels in non-MYCN SHEP1 and SK-N-AS cells (A) and BE(2)-C and LA1-55n cells (B) expressing shGFP or independent shRNAs targeting APRT or HPRT1. mRNA levels were normalized to B2M and are shown relative to shGFP controls.

(C) Immunoblot analysis of HPRT1 in BE(2)-C and SHEP1 cells expressing shGFP or independent HPRT1 shRNAs. PHGDH and SHMT2 are shown for comparison, and α-tubulin was used as a loading control.

(D) Representative phase-contrast images of SHEP1 and SK-N-AS cells expressing shGFP or independent shRNAs targeting APRT or HPRT1. Scale bar, 300 μm.

Data in (A) and (B) are presented as mean ± SD.

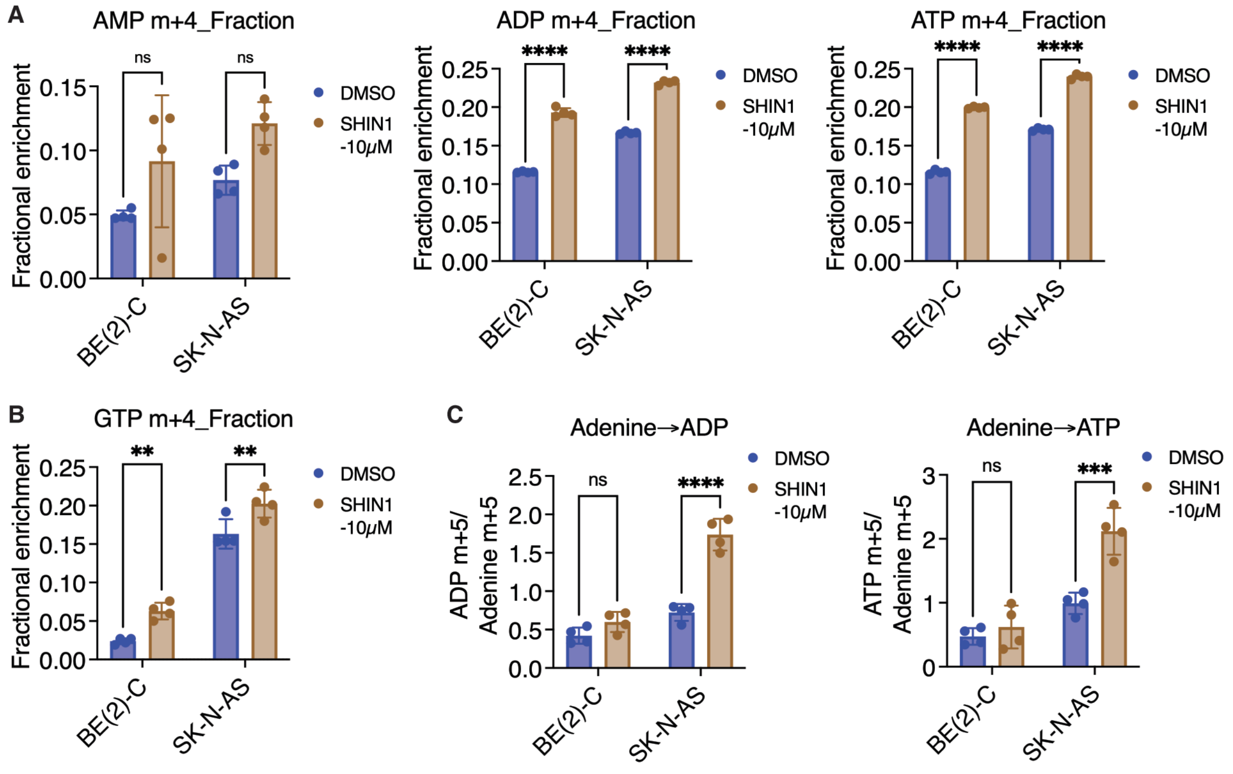


**Figure S5. Effects of SHMT inhibition on hypoxanthine and adenine salvage**

(A-B) MYCN-amplified BE(2)-C and non-MYCN SK-N-AS cells were pretreated with DMSO or SHIN1 (10 μM) for 24 h, followed by 6 h of [$^{15}N_4$]hypoxanthine tracing in the continued presence of DMSO or SHIN1. Fractional enrichment of m+4 AMP, ADP, and ATP (A) and m+4 GTP (B) is shown.

(C) BE(2)-C and SK-N-AS cells were analyzed by [$^{15}N_5$]adenine tracing under the same treatment conditions. Ratios of m+5 ADP and ATP to intracellular [$^{15}N_5$]adenine are shown.

Data are presented as mean ± SD (n = 4). Statistical significance was determined by two-way ANOVA with multiple-comparisons testing. $^{**}P < 0.01$, $^{***}P < 0.001$, $^{****}P < 0.0001$; ns, not significant.

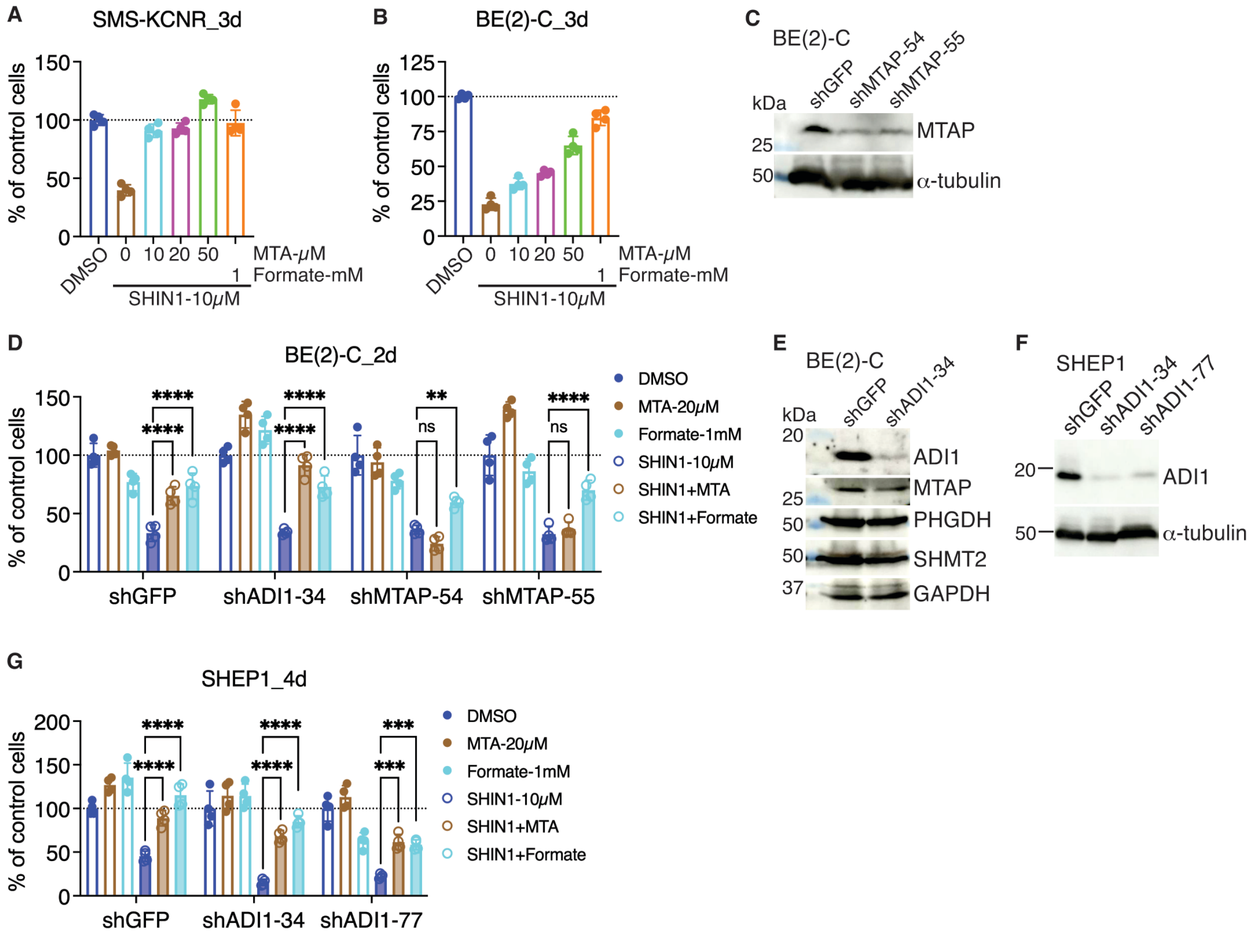


**Figure S6. Methionine salvage supports MTA-mediated rescue through an MTAP-dependent but ADI1-independent mechanism**

(A and B) Growth of MYCN-amplified SMS-KCNR (A) and BE(2)-C (B) cells treated with SHIN1 (10 μM) in the absence or presence of the indicated concentrations of MTA or formate (1 mM) for 3 days. Cell growth is shown relative to DMSO-treated controls.

(C) Immunoblot validation of MTAP knockdown in BE(2)-C cells expressing shGFP or independent MTAP shRNAs. α-Tubulin was used as a loading control.

(D) Growth of BE(2)-C cells expressing shGFP, shADI1-34, shMTAP-54, or shMTAP-55 and treated with DMSO, MTA (20 μM), formate (1 mM), SHIN1 (10 μM), SHIN1 plus MTA, or SHIN1 plus formate for 2 days.

(E) Immunoblot analysis of ADI1, MTAP, PHGDH, and SHMT2 in BE(2)-C cells expressing shGFP or shADI1-34. GAPDH was used as a loading control.

(F) Immunoblot validation of ADI1 knockdown in SHEP1 cells expressing shGFP or independent ADI1 shRNAs. α-Tubulin was used as a loading control.

(G) Growth of SHEP1 cells expressing shGFP, shADI1-34, or shADI1-77 and treated as in (D) for 4 days.

Data are presented as mean ± SD. Statistical significance in (D) and (G) was determined by two-way ANOVA with multiple-comparisons testing. **$P < 0.01$, ***$P < 0.001$, ****$P < 0.0001$; ns, not significant.
